# Short Communication Principal Component Analysis Applied to Alzheimer’s Disease: USA by State

**DOI:** 10.1101/420539

**Authors:** Bodo Parady

## Abstract

Principal Component Analysis (PCA) of Alzheimer’s disease (AD) and twelve epidemiological and socio-economic components of the USA states inform etiology by extracting large scale patterns. The twelve components demonstrate simple pairwise Pearson correlations to AD, and then are analyzed by PCA for loadings associated with AD. Repetitive factor analysis and the culling of questionable data reduced the factors (all per capita) associated with AD to two, one factor with the components dentists and wine consumption and another factor with the components beer consumption and dentists. Dentists and wine are likely associated with reduced AD incidence because of the known inverse association with elevated education. Dental care is known to be inversely associated with AD incidence. The contribution of beer consumption to AD incidence is likely because of the negative effect on the innate immune system from either phytoestrogens in hops, or detritus from fermentation permitting the fungal colonization seen in AD brains.

## 1 Introduction

In 1906 Alois Alzheimer described AD in the eponymous woman Auguste D. [1], [2], and found abnormal deposits of amyloid protein plaques (Aβ) and fibrous protein tangles (neurofibrillary tau). A 1939 review speculated on various causes of AD [3], noting the development of plaques near the site of previous trauma mentioning “intercurrent infection”, suggesting chronic infection is a component of AD. More recently there are a number of reports noting the presence of various fungi in affected portions of all examined AD patient brains [4], [5], [6], [7], [8]. The most common widespread commensal fungus is *Candida albicans* which is in the same family as *Saccharomyces cerevisiae* yeast used in fermentation. This report describes the analysis of the frequency of AD in the states of the USA versus various social and epidemiological factors.

## 2 Methods

To illuminate the etiology of AD, examination of epidemiological data requires a suitable dataset. The WHO dataset was not used because medical identification of AD cases and autopsy procedures lack standardization, and because poverty limits data collection [9]. The USA statewide data is used because of the availability of social and health data, and relatively standard autopsy data is used instead of relying on sometimes subjective AD diagnoses that can conflate vascular dementia and AD. All of the data used in analysis by state, disease, and lifestyle choice is drawn from government and industry public websites and is in Supplemental Table 1 [10], [11], [12], [13], [14], [15], [16], [17], [18], [19], [20].

**Table 1:** Rotated Varimax exploratory component analysis with AD and 12 social and health factors for 46 USA states.

| Factor Matrix (rotated Varimax) | Unweighted, 46 states |  |  |  |  |  |
| --- | --- | --- | --- | --- | --- | --- |
|  | Wealth | Death Rate | Alcohol | AD & Beer | Corr Coef AD | P-value |
| Factor number | 1 | 2 | 3 | 4 |  |  |
| AD deaths per 100k over 65 (AD65) 2014 [11] | 0.256 | -.0085 | 0.197 | 0.824 | 1.000 |  |
| Beer Gal of ethanol per capita 2013 [12] | 0.253 | 0.069 | -0.496 | 0.631 | 0.306 | 0.038 |
| Wine Gal of ethanol per capita 2013 [12] | -0.467 | 0.329 | -0.430 | -0.271 | 0.360 | 0.014 |
| Spirits Gal of ethanol per capita 2013[12] | -0.074 | 0.132 | -0.892 | -0.047 | -0.119 | 0.432 |
| Dentists per 10,000 people 2011 [18] | -0.697 | 0.393 | -0.069 | -0.377 | -0.485 | 0.001 |
| Live births Percent < 2.5 Kg: 2013 [10] | 0.065 | -0.853 | 0.191 | 0.013 | 0.031 | 0.838 |
| Physicians in patient care /10k 2012 [18] | -0.931 | -.0037 | -0.142 | -0.008 | -0.328 | 0.026 |
| Age-adjusted death rate /100k: 2013 [13] | 0.136 | -0.920 | 0.103 | -0.039 | 0.031 | 0.840 |
| Cigarette smoking percent 2014 [14] | 0.518 | -0.643 | -0.054 | 0.200 | 0.323 | 0.028 |
| Smokeless tobacco percent 2010-11 [14] | 0.814 | -0.138 | -0.097 | 0.188 | 0.331 | 0.024 |
| Percent Obese or Overweight 2014 [16] | 0.671 | -0.521 | -0.055 | 0.197 | 0.364 | 0.013 |
| Substance abuse (2010-11) [18] | -0.284 | 0.577 | -0.424 | 0.319 | -0.373 | 0.897 |
| Median per capita income 2014 [19] | -0.718 | 0.224 | -0.373 | -0.235 | -0.373 | 0.011 |
| Sum of squares | 3.7056 | 2.946 | 1.6712 | 1.5669 |  |  |
| Factor loading guidelines (Comrey & Lee, 1992) |  |  |  |  |  |  |
| 1. > .70 - excellent |  |  |  |  |  |  |
| 2. > .63 - very good |  |  |  |  |  |  |
| 3. > .55 - good |  |  |  |  |  |  |
| 4. > .45 - fair |  |  |  |  |  |  |
| 5. > .32 - poor |  |  |  |  |  |  |

Note: Excludes CA, FL, NV, NH

PCA is an established method of multivariate data analysis [21] dating back to Pearson [22] who conceived the notion of a correlation ellipsoid. The idea is to least squares fit low-dimensional subspaces to a cloud of data and relate them to a multidimensional correlation ellipsoid [23]. The analysis goal is to reduce the linear combinations of covariates that are uncorrelated and have maximal variance, i.e. have axes at perpendicular maxima on the correlation ellipsoids.

Epidemiological, social and health data from all states are analyzed by creating a correlation matrix, and from this correlation matrix, PCA and Varimax rotation are applied for exploratory component analysis [24] [25]. Factor loading levels are evaluated according to standard practice [26]. The state data for the items in the cross-correlation matrix are examined for unweighted correlation coefficients [27] measured against AD65 (AD cases per 100,000 of the over 65 population) assuming that all of the socio-economic factors apply evenly over a lifetime. Factors were removed when the communal loadings were too low to warrant including them in a 4 factor analysis. Data from CA, FL, NV, and NH were excluded for various reasons. Nevada was removed because it has a many celebratory visitors that alter alcohol consumption, and similarly New Hampshire has no sales tax, driving significant cross border alcohol sales [28]. Florida and California were excluded because of the large differences between the elderly social profiles and the youth social profiles. The medium and small states tend to have lowest mobility, and highest proportion of older residents living most of their lives in one state. In the remaining 46 state data there are correlations between AD65 and beer consumption, wine consumption, dentists per capita, doctors per capita, cigarette smoking, smokeless tobacco, overweight percentage, and income level.

## 3 Results

AD65 and epidemiological and socio-economic data from the 46 states are calculated into a correlation matrix (Supplemental Table 1). Varimax factor analysis is then applied and is shown in Table 1. Four factors account for 76% of the variance in the data.

Factor analysis shows that AD65 and per capita beer consumption are the most significant components of Factor 4 in Table 1, and they are positively correlated (r=0.306, p=0.038). None of the other loadings in Factor 4 were significant although there were minor loadings for dentists per capita and substance abuse. Thus 6.35% of the variance in the state data is attributed to AD65 or beer consumption, and neither appear as significant loadings in the other factors. 43% of the variance in the socio-economic data is attributed to Factor 1, which is composed of various economic components that refer to the wealth of the various states. Factor 2 loadings are labeled “Death Rate”, and may be summarized as “risky and poor life choice behavior leading to early death and thus increasing age adjusted death rate.” Cigarette smoking, overweight and substance abuse are major components of Factor 2. Altogether Factor 2 contributes 16% of the total variance and AD65 has no loading it. The use of AD65 instead of age adjusted death rate (AADR) due to AD is validated by AD65 showing no loading for Factor 2: Since AD can only occur at maturity, and because early death reduces the occurrence rate of AD, use of AADR due to AD induces a confounding anti-correlation with overall AADR. Factor 3, labeled “Alcohol”, includes significant loadings from all forms of alcohol consumption and addictive behavior (9.4% of the variance), and again, no loadings are associated with AD65.

Starting with this initial exploratory PCA, successive reductions were performed on the cofactors of AD65 to remove components with high common loadings: e.g. physicians per capita and retaining dentists per capita both have high loadings in the primary wealth factor, and additionally dentists have some loading in the AD/Beer factor which makes Physicians per capita a redundant wealth factor, lacking any other loadings. Also, beer consumption in gallons of beer per capita replaced gallons of ethanol in beer in final analysis because a non-alcohol component appears associated with the incidence of AD; this change increases Pearson correlation and decreases p.

Washington state data was removed from analysis because of a change in reporting procedure before 2000 that exaggerates the diagnosis of AD at death. For every year since 2000 Washington is the state with the highest annual AD65 (e.g. Supplemental Figure 1), and has a raw AD death rate and AD AADR of approximately 44 per 100,000 population up from 11 per 100,000 before 1998. Removing the Washington state data from analysis elevated all correlation coefficients and reduced all p values for wine, beer and dentists per capita, suggesting anomaly.

The results of component reduction are shown in Table 2. Wealth/Education is labeled that way, because wine consumption is seen here as a proxy for education [30] and wealth. Beer has no loading in the Wealth/Education factor and dominates factor 2. AD65 loading is evenly split between the two factors which means that AD incidence likely has two cofactors in this analysis: One is wealth and the other is associated with beer and dentistry, but neither wine nor spirits. Dentist loadings have two aspects: Dentists are associated with a wealthy population, and they also manage the oral hygiene of the population. Better dental hygiene has been associated with reduction and management of fungal infections and biofilms [31] [32] [33] which could enter the brain through oropharyngeal tracts. The components of factor 2 point to a biological, not socio-economic contribution.

**Table 2:**
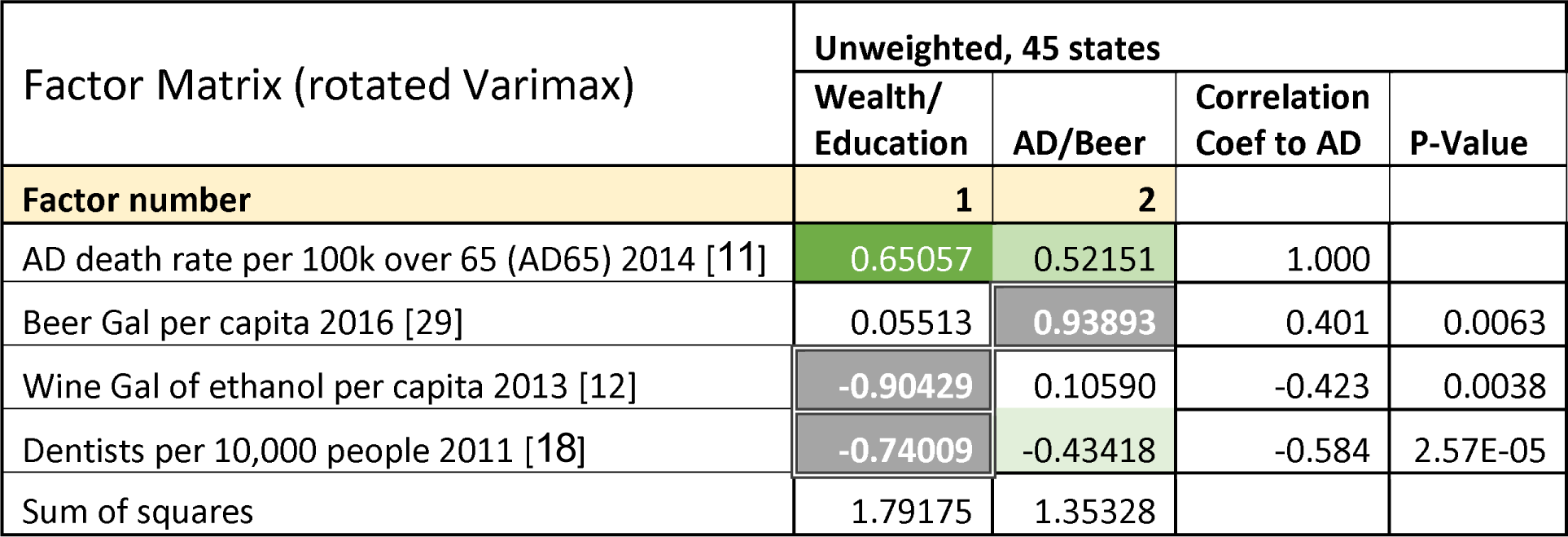
Reduced rotated Varimax confirmatory component analysis with AD, beer, wine and dentists as factors for 45 USA states (excludes Washington).

| Factor Matrix (rotated Varimax) | Unweighted, 45 states |  |  |  |
| --- | --- | --- | --- | --- |
|  | Wealth/<br>Education | AD/Beer | Correlation<br>Coef to AD | P-Value |
| Factor number | 1 | 2 |  |  |
| AD death rate per 100k over 65 (AD65) 2014 [11] | 0.65057 | 0.52151 | 1.000 |  |
| Beer Gal per capita 2016 [29] | 0.05513 | 0.93893 | 0.401 | 0.0063 |
| Wine Gal of ethanol per capita 2013 [12] | -0.90429 | 0.10590 | -0.423 | 0.0038 |
| Dentists per 10,000 people 2011 [18] | -0.74009 | -0.43418 | -0.584 | 2.57E-05 |
| Sum of squares | 1.79175 | 1.35328 |  |  |

## 4 Discussion

High Pearson correlations are found for per capita wine and beer consumption and per capita dentists compared to AD65. Of the initial 12 component study, only state-wise per capita beer ethanol consumption has strong loading with AD65 in PCA Varimax rotation. Because of the highly correlated factors within wealth/poverty, age adjusted death rates, and risky behavior, PCA is used to isolate collinearity and identify confounding multivariate relationships. PCA deals with the problem of multi-collinearity which otherwise leads to instabilities in multivariate regression analysis [34]. Factor analysis reduced the number of linear covariates to AD65 and verified that AD65 had no extraneous loading with age adjusted death rates (Table 1).

Lower education level is a well-known risk factor of AD incidence [35] [36] and wine [37] [38] is well correlated to elevated wealth and education [39]. The frequent comment when beer is shown correlated with AD is: “Beer drinking is probably associated with class or poverty.” The loadings in factor analysis show that beer consumption factor loadings with AD65 are independent of the wealth/poverty factor.

Published work indicates that beer might be a factor in the development of AD and dementia [40]. Iceland has lowest rate of AD in Europe with 1.19% of the population which is much lower than the EU average AD rate of 1.55% [41]. A fact that supports a role for beer in the development of AD is that Iceland banned beer until 1989 [42]. Another factor in the development of AD could be the extremely low incidence of mild hyperthyroidism in Iceland (4x lower) compared to the incidence of mild hyperthyroidism in Europe’s elderly [43]. Thyroid disease was identified as a risk factor for AD in a Canadian study [44]. Beer consumption (>1000ml/week) was associated with goiter incidence in a German study [45] which suggests that beer interferes with iodine mobilization. Note the first case of AD was in Auguste D. from the famous beer consuming region of Bavaria [11].

The Copenhagen City Heart study found that the relative risk (RR) of all dementia was around 2x for occasional to daily beer drinkers [46]. RR was 2.47 for light and moderate beer drinkers in a multi-city study in China [47]. And RR was 1.96 for daily beer drinkers in a New England study [48]. The Helsinki study of beer drinkers at autopsy found reduced Aβ aggregation [49], but they cautioned that the study was too small to observe relative risk of AD onset; reduction of Aβ aggregation may be evidence of innate immune suppression.

Of the reasons to associate beer consumption with the development of AD, consider either yeast (live or dead and its detritus), or hops. Yeast might overload the innate immune system and limit control of systemic fungal/yeast infections [50]. Hops contain high levels of phytoestrogens which may induce many estrogenic disorders [51], including “brewer’s droop [52].” Women with nM levels of estrogen/estradiol have the highest risk of AD (those with 0 or µM levels have lowest risk of AD) [53]. These nM levels may correspond to the significant increases in phytoestrogens from hops [54], and estradiol from drinking beer or wine [55]; after consuming beer (alcoholic or non-alcoholic) the estradiol levels in men appear to be almost 3x those found in postmenopausal women.

Results here are in line with smaller studies that find reduced risk for AD associated with both wine (an education correlate) and the availability of dental care (related to wealth and the risk factors associated with beer), and elevated risk of AD for beer consumption irrespective of wealth. There is a need to establish uniform reporting rules for AD at death to make USA data more consistent for epidemiological studies.

## 5 Acknowledgements

The author wishes to thank Prof Bruce Ames for encouragement, and Luis Carrasco Llamas for offering updated research findings.

## 6 Supplement

**Supplemental Figure 1:**
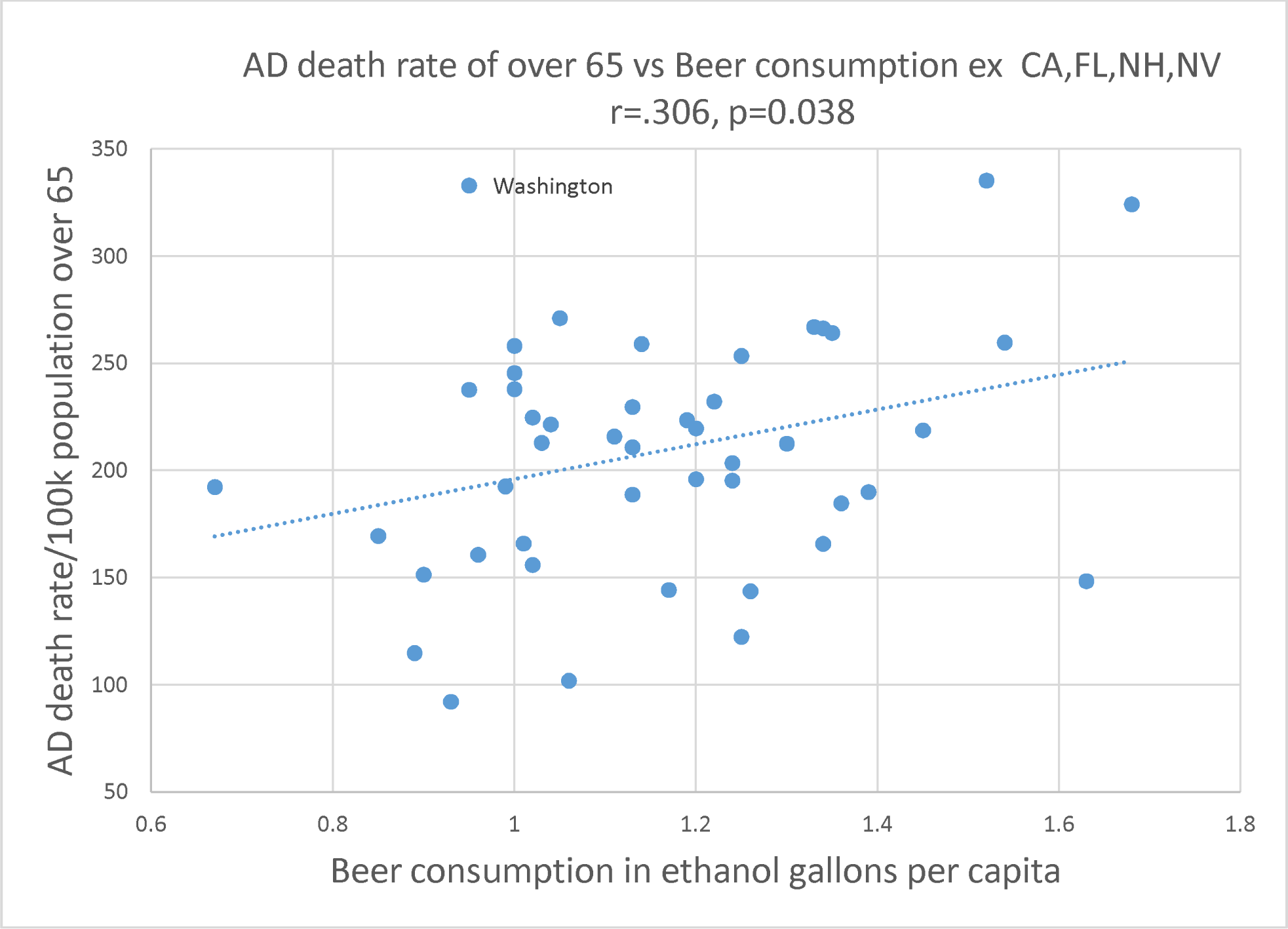
46 State Per capita beer consumption and age adjusted AD mortality rate for 2014.

**Supplemental Table 1:** 46 state epidemiological and socio-economic data.

| State | AD death rate per 100k over 65 2014 | Beer Gal per capita 2013 | Wine Gal per capita 2013 | Spirits Gal per capita 2013 | Dentists per 10,000 people 2011 | Live births Percent less than 2.5 Kg: 2013 | Physicians in patient care /10,000 2012 | Age-adjusted death rate per 100k: 2013 | Cigarette smoking percent 2014 | Smoke less tobacco percent 2010-11 | Percent Obese or Over weight 2014 | Substance abuse 2010-11 | Median per capita income 2014 |
| --- | --- | --- | --- | --- | --- | --- | --- | --- | --- | --- | --- | --- | --- |
| Alabama | 259.024 | 1.14 | 0.25 | 0.6 | 4.4 | 9.98 | 20.5 | 1037.9 | 21.1 | 6.1 | 67 | 0.067 | \$23,606 |
| Alaska | 101.936 | 1.06 | 0.52 | 1.16 | 7.4 | 5.81 | 22.6 | 944.6 | 19.9 | 6.8 | 64.8 | 0.093 | \$33,062 |
| Arizona | 229.646 | 1.13 | 0.42 | 0.81 | 5.1 | 6.95 | 21 | 873.5 | 16.5 | 3.2 | 64 | 0.103 | \$25,715 |
| Arkansas | 258.144 | 1 | 0.2 | 0.6 | 3.9 | 8.84 | 19.4 | 996.3 | 24.7 | 6.9 | 70.6 | 0.068 | \$22,883 |
| Colorado | 195.977 | 1.2 | 0.5 | 1.11 | 6.5 | 8.77 | 25 | 856.1 | 15.7 | 4.2 | 57.4 | 0.099 | \$32,357 |
| Connecticut | 169.471 | 0.85 | 0.61 | 0.91 | 7.5 | 7.8 | 33.9 | 857.5 | 15.4 | 1.8 | 60.4 | 0.09 | \$39,373 |
| Delaware | 122.418 | 1.25 | 0.72 | 1.63 | 4.6 | 8.32 | 24.5 | 1001.9 | 19.9 | 2.2 | 67.4 | 0.079 | \$30,488 |
| Georgia | 212.912 | 1.03 | 0.25 | 0.69 | 4.4 | 9.36 | 19.9 | 1037.4 | 29.2 | 5 | 65.7 | 0.069 | \$25,615 |
| Hawaii | 143.695 | 1.26 | 0.54 | 0.78 | 8.1 | 8.19 | 29.7 | 752.2 | 14.1 | 1.7 | 58.1 | 0.086 | \$29,736 |
| Idaho | 160.729 | 0.96 | 1.08 | 0.77 | 5.8 | 6.47 | 17.4 | 856.6 | 15.9 | 5.7 | 65.7 | 0.092 | \$23,938 |
| Illinois | 188.774 | 1.13 | 0.42 | 0.78 | 6.3 | 8.19 | 26.1 | 973.8 | 16.5 | 2.6 | 63.8 | 0.084 | \$30,417 |
| Indiana | 237.963 | 1 | 0.28 | 0.81 | 4.7 | 7.97 | 21 | 962 | 22.9 | 4.9 | 66.5 | 0.083 | \$25,140 |
| Iowa | 266.964 | 1.33 | 0.22 | 0.8 | 5.3 | 6.58 | 19.7 | 848.2 | 18.5 | 4.9 | 66.9 | 0.087 | \$28,361 |
| Kansas | 192.595 | 0.99 | 0.14 | 0.76 | 5 | 7.12 | 22.4 | 867.2 | 18.1 | 5.5 | 66 | 0.084 | \$27,870 |
| Kentucky | 237.669 | 0.95 | 0.2 | 0.7 | 5.6 | 8.82 | 22 | 1024.5 | 26.2 | 7 | 66.7 | 0.064 | \$23,684 |
| Louisiana | 266.353 | 1.34 | 0.32 | 0.93 | 4.7 | 10.88 | 24.3 | 1074.6 | 24 | 5.7 | 68.9 | 0.075 | \$24,800 |
| Maine | 184.710 | 1.36 | 0.42 | 0.91 | 5 | 6.82 | 28.7 | 918.7 | 19.3 | 2 | 64.5 | 0.068 | \$27,978 |
| Maryland | 114.858 | 0.89 | 0.39 | 0.9 | 7.3 | 8.71 | 35.4 | 985.2 | 14.6 | 2.5 | 64.9 | 0.068 | \$36,338 |
| Massachusetts | 165.945 | 1.01 | 0.66 | 0.9 | 8.4 | 7.63 | 39.6 | 884.8 | 14.7 | 1.5 | 58.9 | 0.102 | \$36,593 |
| Michigan | 224.725 | 1.02 | 0.37 | 0.85 | 6.1 | 8.34 | 26 | 966 | 21.2 | 3.9 | 65.6 | 0.086 | \$26,613 |
| Minnesota | 210.859 | 1.13 | 0.43 | 1.16 | 6.1 | 6.47 | 27.2 | 825.2 | 16.3 | 5 | 64.1 | 0.091 | \$32,638 |
| Mississippi | 264.180 | 1.35 | 0.16 | 0.73 | 3.9 | 11.67 | 17.3 | 1071.4 | 23 | 8.5 | 70.7 | 0.071 | \$21,036 |
| Missouri | 223.516 | 1.19 | 0.36 | 0.89 | 4.7 | 7.89 | 24.1 | 952.4 | 20.6 | 5.1 | 65.6 | 0.073 | \$26,126 |
| Montana | 148.405 | 1.63 | 0.49 | 0.95 | 5.7 | 7.31 | 21.9 | 890.2 | 19.9 | 8 | 63 | 0.103 | \$25,989 |
| Nebraska | 189.975 | 1.39 | 0.21 | 0.72 | 6.2 | 6.58 | 23.5 | 867.9 | 17.3 | 5.3 | 66.7 | 0.08 | \$27,446 |
| New Jersey | 151.450 | 0.9 | 0.62 | 1.03 | 8 | 8.32 | 30.1 | 956 | 15.1 | 1.7 | 63.1 | 0.082 | \$37,288 |
| New Mexico | 144.267 | 1.17 | 0.34 | 0.85 | 4.6 | 8.82 | 22.2 | 891.9 | 19.1 | 4.3 | 64.9 | 0.092 | \$23,683 |
| New York | 92.140 | 0.93 | 0.51 | 0.77 | 7.7 | 8 | 35 | 973.7 | 14.4 | 2.2 | 61.1 | 0.084 | \$33,095 |
| North Carolina | 221.508 | 1.04 | 0.39 | 0.6 | 4.5 | 8.87 | 23.4 | 986 | 19.1 | 4.3 | 65.6 | 0.07 | \$25,774 |
| North Dakota | 324.191 | 1.68 | 0.32 | 1.37 | 5.1 | 6.42 | 24.1 | 818.4 | 19.9 | 7.6 | 68.8 | 0.094 | \$33,071 |
| Ohio | 232.175 | 1.22 | 0.3 | 0.53 | 5.2 | 8.57 | 26.2 | 967.4 | 21 | 4.2 | 66.7 | 0.089 | \$26,937 |
| Oklahoma | 215.871 | 1.11 | 0.19 | 0.63 | 5 | 8.17 | 19.3 | 961.4 | 21.1 | 6.3 | 68.2 | 0.089 | \$25,229 |
| Oregon | 219.645 | 1.2 | 0.57 | 0.89 | 6.8 | 6.19 | 26.4 | 893 | 17 | 4.6 | 61.7 | 0.098 | \$27,646 |
| Pennsylvania | 165.805 | 1.34 | 0.31 | 0.68 | 6.2 | 8.09 | 29.6 | 963.4 | 11.3 | 4.4 | 64.1 | 0.089 | \$29,220 |
| Rhode Island | 245.496 | 1 | 0.58 | 0.98 | 5.5 | 7.46 | 34.6 | 889.6 | 16.3 | 1.9 | 62.4 | 0.107 | \$30,830 |
| South Carolina | 253.473 | 1.25 | 0.25 | 0.81 | 4.6 | 9.7 | 21.6 | 1030 | 21.5 | 4.4 | 67 | 0.081 | \$24,596 |
| South Dakota | 335.262 | 1.52 | 0.28 | 1.01 | 5 | 6.24 | 22.2 | 846.4 | 18.6 | 6.6 | 65.2 | 0.1 | \$26,959 |
| Tennessee | 271.081 | 1.05 | 0.25 | 0.69 | 4.9 | 9.11 | 24.8 | 1011.8 | 24.2 | 4.8 | 67.1 | 0.082 | \$24,922 |
| Texas | 212.534 | 1.3 | 0.32 | 0.63 | 4.5 | 8.35 | 20.3 | 947.6 | 14.5 | 4.3 | 67.8 | 0.08 | \$27,125 |
| Utah | 192.309 | 0.67 | 0.19 | 0.51 | 6.4 | 6.92 | 19.6 | 823.2 | 9.7 | 2.9 | 59.5 | 0.063 | \$24,877 |
| Vermont | 259.682 | 1.54 | 0.74 | 0.72 | 5.8 | 6.51 | 33.3 | 908.6 | 16.4 | 2.8 | 60.2 | 0.1 | \$29,178 |
| Virginia | 155.971 | 1.02 | 0.46 | 0.62 | 6 | 8.04 | 25.8 | 963.1 | 19.5 | 4 | 64.7 | 0.082 | \$34,052 |
| Washington | 332.961 | 0.95 | 0.53 | 0.76 | 7 | 6.23 | 25.1 | 869.4 | 15.3 | 3.7 | 63.4 | 0.087 | \$31,841 |
| West Virginia | 195.316 | 1.24 | 0.1 | 0.46 | 4.7 | 9.4 | 23.8 | 1031.5 | 26.7 | 9.4 | 69.6 | 0.071 | \$22,714 |
| Wisconsin | 218.729 | 1.45 | 0.38 | 1.18 | 5.7 | 7.12 | 24.9 | 879.1 | 17.4 | 4.3 | 67.4 | 0.086 | \$28,213 |
| Wyoming | 203.489 | 1.24 | 0.29 | 1.13 | 5 | 8.43 | 18.8 | 897.4 | 19.5 | 8.8 | 64.6 | 0.083 | \$29,698 |

